# Functional MRI based simulations of ECoG grid configurations for optimal measurement of spatially distributed hand-gesture information

**DOI:** 10.1101/2020.12.27.424363

**Authors:** Max van den Boom, Kai J. Miller, Nick F. Ramsey, Dora Hermes

## Abstract

In electrocorticography (ECoG), the physical characteristics of the electrode grid determine which aspect of the neurophysiology is measured. For particular cases, the ECoG grid may be tailored to capture specific features, such as in the development and use of brain-computer-interfaces (BCI). Neural representations of hand movement are increasingly used to control ECoG based BCIs. However, it remains unclear which grid configurations are the most optimal to capture the dynamics of hand gesture information. Here, we investigate how the design and surgical placement of grids would affect the usability of ECoG measurements. High resolution 7T functional MRI was used as a proxy for neural activity in ten healthy participants to simulate various grid configurations, and evaluated the performance of each configuration for decoding hand gestures. The grid configurations varied in number of electrodes, electrode distance and electrode size. Optimal decoding of hand gestures occurred in grid configurations with a higher number of densely-packed, large-size, electrodes up to a grid of ^~^5×5 electrodes. When restricting the grid placement to a highly informative region of primary sensorimotor cortex, optimal parameters converged to about 3×3 electrodes, an inter-electrode distance of 8mm, and an electrode size of 3mm radius (performing at ^~^70% 3-class classification accuracy). Our approach might be used to identify the most informative region, find the optimal grid configuration and assist in positioning of the grid to achieve high BCI performance for the decoding of hand-gestures prior to surgical implantation.

## Introduction

The human brain can be explored at many different spatial scales, from (sub)millimeter to centimeter resolution. Each scale of measurement provides an unique window into the underlying neuronal activity (Freeman 2000). As a result, research is often bound by the resolution at which information is captured. An empirical understanding of which resolution provides the most information for the interpretation of a particular brain function is essential for the development of brain-computer interfaces (BCI).

Electrocorticography (ECoG-) based BCIs utilize properties of intracranial electrode grids to record the electrical field potential from neuronal populations (Crone et al. 1998; Ramsey et al. 2006; Schalk et al. 2008; Miller et al. 2010; Yanagisawa et al. 2012; Vansteensel et al. 2016; Benabid et al. 2019). The ECoG signals are translated or interpreted to understand brain activity or to control external devices. ECoG grids can be configured in a number of ways: spanning small or large areas of cortex with different electrode sizes and densities. Currently, several different grid configurations are used in research, varying from traditional clinical electrode grids and strips with 1 cm inter-electrode distance to high-density research grids (Buzsáki et al. 2012). The configuration of an ECoG electrode grid determines the resolution and scale with which the underlying neuronal activity is measured, and therefore which information is conveyed to a BCI. Therefore, it is essential to know which configuration provides the most useful information for control. In practice, comparing different electrode configurations on the same brain region in human subjects is not possible given the invasive nature of ECoG recordings and the fact that this is done in a clinical setting. Given the limitations in experimental testing and the need for regulatory approval of implanted BCI systems in humans (Vansteensel et al. 2016; Benabid et al. 2019; Miller et al. 2020), it is essential to understand which scale provides most information about brain functions used in BCIs.

Hand-representations are a promising target for use in BCIs (Bleichner et al. 2014, 2016; Brandman et al. 2017). During hand movement, a robust neurophysiological response occurs in the contralateral sensorimotor cortex (Crone et al. 1998; Miller et al. 2007) with a focal increase of high-frequency band (HFB) power and a more distributed decrease in low-frequency band power (Hermes et al. 2012). Subsequent ECoG research showed that the representations of different fingers can be distinguished in the primary sensorimotor cortex using a HFB component (Miller et al. 2009). Given the increased interest in the use of hand-representations in BCI solutions (Pistohl et al. 2012; Chestek et al. 2013; Collinger et al. 2013; Bleichner et al. 2016; Vansteensel et al. 2016; Brandman et al. 2017) it is essential to understand which grid configuration would optimally capture hand movement information. Recent research has shown that the ECoG HFB signal correlates well with the BOLD response at the standard clinical scale (Hermes et al. 2012), that the millimeter scale finger representations found in ECoG HFB activity are well matched with 7T fMRI (Siero et al. 2014) and that even non-linearities in 7T fMRI match the ECoG HFB responses (Siero et al. 2013). Therefore, even though fMRI is a less direct measure of neuronal activity than electrode recordings, the two modalities do correlate well across multiple spatial scales, suggesting that we can leverage 7T fMRI to simulate different grid configurations and estimate the electrode scale and density that would provide the most information to decode hand-movement.

In this study, we specifically investigate which ECoG scale captures the most information about hand-gestures. Using 7T fMRI measurements, we simulated the placement of different grid configurations on the exposed brain surface, varying the number of electrodes, inter-electrode (center-to-center) distance and size in a similar manner to variations found in commercially-available ECoG implants.

## Methods

### Participants

Ten healthy subjects (age 23.9 ± 5.1 years, 6 female) participated in the experiment. All participants had normal or corrected-to-normal vision. The study was approved by the ethics committee of the University Medical Center Utrecht (ref 13-585) and participants gave written informed consent to participate, in accordance with the Declaration of Helsinki (2013). According to the Edinburgh Inventory (Oldfield, 1971), all were right-handed.

### Task

Participants were asked to execute three hand gestures taken from the American Sign Language alphabet (the letters S, F, and L), shown in Figure 1. The gestures were chosen to maximize the difference between flexion and extension of the finger combinations. All participants were naïve to sign language and were trained prior to the experiment on the three letter-gesture combinations. While in the scanner, one of the three letters was presented each trial. Each letter was shown for either 3, 5 or 7 seconds, followed by a fixation cross shown for 12 seconds. Participants were instructed to perform the corresponding gesture upon appearance of the letter and hold the gesture until the letter disappeared. A total of 45 letters were presented at random, ensuring 15 trials per letter. The different durations were equally balanced over the letter conditions.

**Figure 1.**
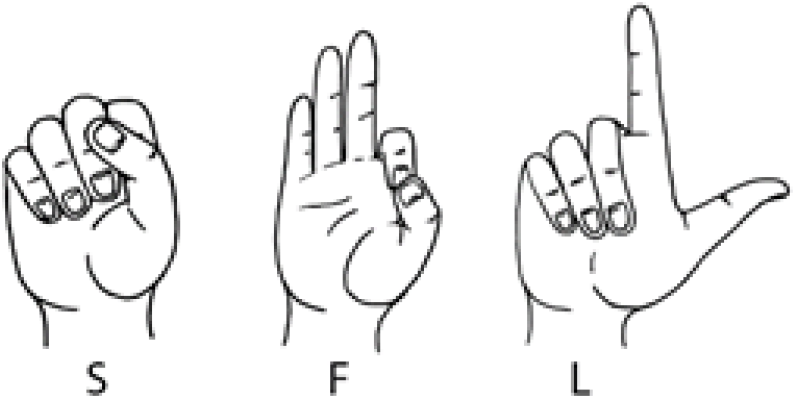
Hand gestures used in this study. The three hand gestures were taken from the American Sign Language alphabet and involved the S, F and L. Note that during the task only a letter was presented, while the participant performed the corresponding gesture.

### MRI data acquisition

MRI data were acquired using a 7T Philips Achieva MRI system with a 32-channel head coil. Functional MRI data were recorded using an EPI sequence (TR/TE = 1600/27ms, FA = 70 degrees, 26 slices centered around the motor cortex, acquisition matrix 144 x 144, slice thickness 1.60 mm with no gap, 1.47 mm in plane resolution); shared datasets can be found on OSF: https://osf.io/z6j3x/. A T1-weighted image (TR/TE: 7/3.05ms; FA: 8; resolution 0.78 x 0.78 mm in-plane, slice thickness 0.8mm, no gap) was acquired for anatomical reference. The cues were visually projected onto a mirror attached to the head coil. Each participant performed a single functional run of the task with 481 volumes (5 dummy volumes) which took about 13 minutes.

### Preprocessing

Preprocessing of the imaging data was performed with the use of SPM12, FSL and custom MATLAB (Mathworks Inc.) scripts. The anatomical image was skull-stripped using the white and grey matter segmentation output and converted to a 3D brain surface (Dale et al. 1999) using Freesurfer (http://surfer.nmr.mgh.harvard.edu/). Functional images were slice-time corrected, realigned to correct for motion and corrected for geometric distortions (Andersson et al. 2003). The corrected functional images were aligned to the anatomical scan by transformation of their affine matrices. No reslicing/resampling was done at this stage to prevent a potential loss of decodable information.

### Grid simulation

In order to investigate which different ECoG grid configurations would capture hand-gesture information best, we simulated different grid configurations on the exposed (unwarped) brain surface convexity to assess how different grid configuration parameters influence decoding performance of the three gestures from fMRI data. To achieve the most realistic simulations, all analyses were done in the subject’s native space. The process of simulation involved three steps: positioning the centers of the virtual grids (Figure 2A), placement of different grid configurations around this center (Figure 2B) and the decoding of gestures based on the underlying voxels from the functional volumes to evaluate the information that is captured by each grid (Figure 2C).

**Figure 2.**
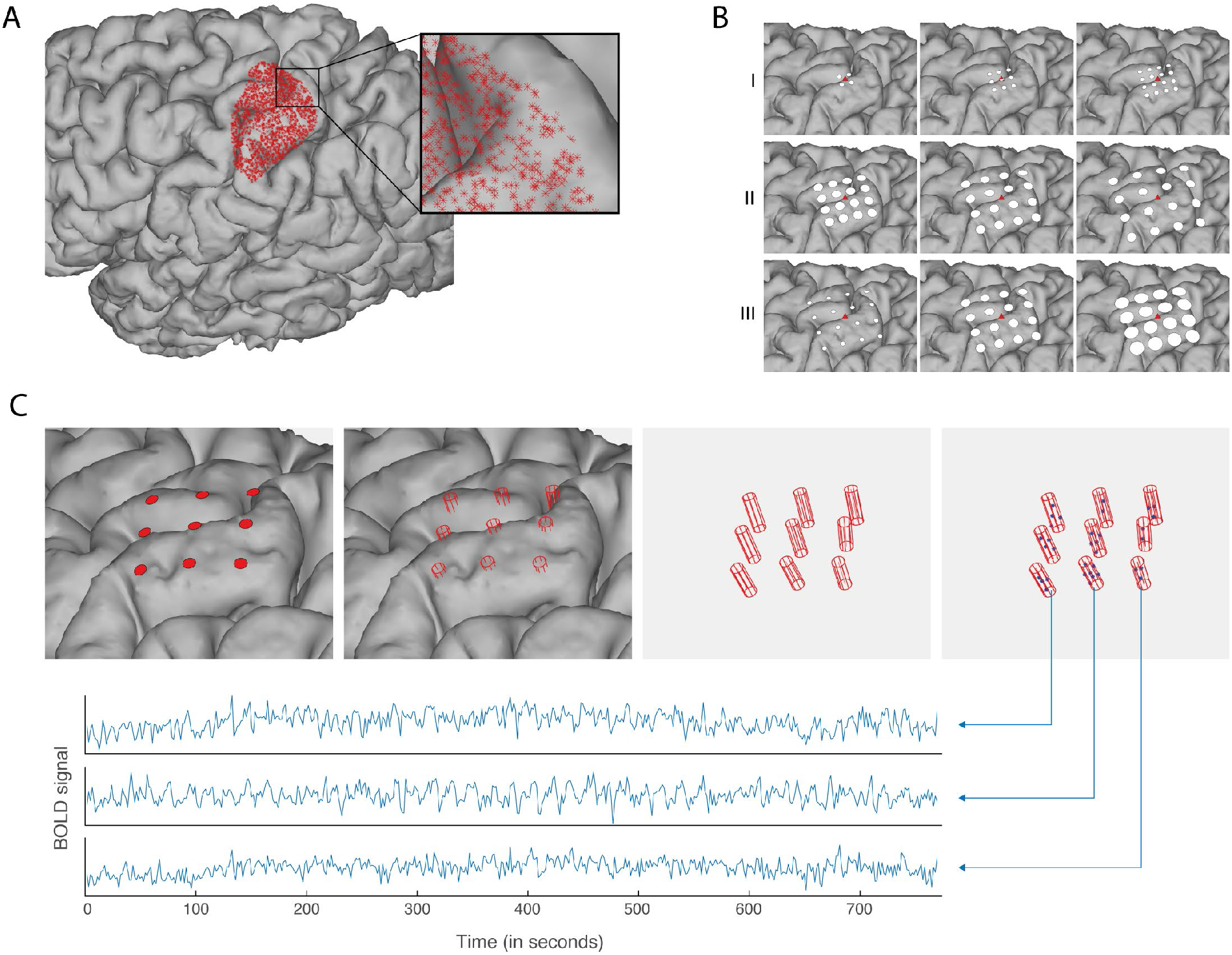
The grid simulation procedure (Subject 7). (A). The centers of the virtual grids were positioned over an empirically-identified cortical hand-gesture area. A searchlight classification evaluated 200,000 potential center positions for underlying hand-gesture information. Positions that yielded classification above the 95% percentile were kept (^~^17,000 positions). A subset of 2,000 positions were selected at random and served as the centers for the simulation of different grid configurations. (B). Different grid configurations were simulated at each of the 2,000 positions. The center of each simulated grid is shown as a red triangle, the white discs indicate the placement of virtual electrodes given different grid configurations. Grid configurations varied in number of electrodes (2×2… 8×8), inter-electrode distance (4mm, 6mm, 8mm, 10mm center-to-center) and electrode size (radius of 1mm, 2mm, 3mm). A total of 84 grid configurations were simulated per position. (C). All of the 2,000 center positions, with each 84 grid placements, were evaluated by the classification of the hand-gestures. Classification was performed using a support vector machine, where each electrode represents one input feature. The signal for each electrode was the average BOLD amplitude of the gray matter voxels in the functional volumes that fell inside of the electrode cylinder.

The centers of the grids were positioned over an area in the brain that has meaningful information related to the hand motor task. In order to find that meaningful area, a convex brain surface hull was produced for each participant, covering the dorsal and lateral side of the frontal-and parietal lobe. Next, 200,000 potential grid positions were generated on the hull at random. Decoding performance for each position was evaluated by performing multivariate searchlight classification (Kriegeskorte et al. 2006) using the voxels in the functional volumes within a 7mm radius as features to classify on. A searchlight approach was preferred over the use of activity amplitude as it provides a more qualitative estimation of the local information (Kriegeskorte and Bandettini 2007). After the classification of all potential positions, only the positions with a classification accuracy score above the 95^th^ percentile were kept and used as centers for the simulated grids. For every participant, this procedure resulted in a cluster of positions over the sensorimotor cortex roughly overlapping the hand-knob area, shown in Supplement Figure 1. For five participants, several smaller clusters of positions were identified in addition to the large cluster around primary sensorimotor areas. In such cases, only the largest sensorimotor cluster was kept. On average, a sensorimotor cluster contained ^~^17000 positions. To limit the number of calculations, 2,000 positions were picked at random for grid simulation (Figure 2A).

During grid placement, each of the 2,000 positions received a random rotation and served as the center for multiple grid projections, as demonstrated for a single position in figure 2B. Grid configurations varied in number of electrodes (2×2, 3×3, 4×4, 5×5, 6×6, 7×7, 8×8), inter-electrode distance (4mm, 6mm, 8mm and 10mm center-to-center) and electrode size (radius 1mm, 2mm, 3mm). As a result, a total of 84 (7×4×3) virtual grids were “placed” per position.

After placement, each grid was used to classify the three executed hand gestures by taking the signal from the grey matter fMRI voxels underneath each virtual grid electrode, shown in Figure 2C. For each electrode, a cylinder with the radius of the electrode was projected perpendicular to the convex hull of the cortical surface reaching 5mm downward and 5mm upward. Only the gray matter voxels (according to the Freesurfer segmentation) from the functional volumes within each electrode cylinder were used for classification. Because the virtual electrodes were positioned onto a hull of the brain, it is possible that they fell slightly inside of the gyri; The inclusion of voxels upward from the electrode made certain that no gray matter voxels were excluded, which in reality would be underneath the electrodes. We required a minimum of two features (i.e. electrodes) in each grid configuration. In addition, to prevent grids with a large surface (i.e. high number of electrodes and/or inter-electrode distance) from covering and classifying from regions far outside of the primary sensorimotor, only voxels within 15mm of the sensorimotor cortex (as defined by the Desikan-Killiany atlas; Desikan et al. 2006) were used during classification. The MATLAB scripts that were used to implement the grid simulations are publicly available through the open science framework (OSF: https://osf.io/z6j3x/).

### Classification procedure

The classification procedure was implemented using custom MATLAB (Mathworks Inc.) code provided alongside this article (OSF: https://osf.io/z6j3x/). Before classification, the raw BOLD signal of each voxel was normalized (division by its mean and multiplication by 100) to reflect the percentage signal change, and detrended (linear, quadratic and cubic). Classification was performed using a Support Vector Machine (SVM) and a linear kernel (Bishop 2006).

The voxels or combination of voxels that were used as features in the SVM differed per step of the simulation procedure. During the step of grid positioning, a searchlight was used, where each grey matter voxel within the searchlight served as a feature in the SVM. During the step of decoding grids, the gray matter voxels within each virtual electrode (i.e. cylinder) were averaged and each electrode average served as a feature in the SVM.

The BOLD response that we used in the SVM was determined as follows. For each trial, we took the time-period around the expected peak of the hemodynamic response (*t_startHRF_-t_endHRF_*) based on the trial’s duration (3, 5, or 7 seconds). Within that time-period, we calculated the average over all features *<u>X</u>*(*t*)and determined the time *t_max_* that yielded the highest BOLD peak (where *<u>X</u>*(*t*)was maximum). The BOLD signal of each virtual electrode at that scan time *<u>X</u>*(*t_max_*)served as a feature in the classification. Multi-class classification (3 gestures) was achieved by applying a one-versus-all classification scheme, where every class is classified against the data of all other classes together, and the winner (furthest from the hyperplane) takes all. The decodability of the three gestures was determined using a leave-one out cross validation and is expressed as a classification accuracy score, which is the percentage of trials that were predicted correctly.

### Two ways to position grids

In the first analyses, we placed the center of the simulated grid in the area with meaningful information about the gestures (Figure 2A). However, as the number of simulated electrodes increases, grids become larger than the informative region and extend beyond it. To investigate whether classification could be optimized by restricting all electrodes in the stimulated grids to the informative region, we also ran the simulations while limiting the grids to entirely fall within the region of interest indicated by the searchlight.

### Statistics

To determine whether a classification score was significantly above chance, we used the cumulative binomial distribution. Given 45 trials, a chance level of 33% and a cumulative probability of 95% (one-sided, α = .05), classification scores above 44% were considered significant.

The effects of the variations in grid configuration were evaluated by a full factorial analysis of variance (ANOVA). For each of the 10 participants and each grid configuration (7×4×3 = 84), the average over the 2,000 classification accuracy scores was used as the dependent variable. The ANOVA evaluated the effects of three properties (as independent variables): the number of electrodes (7 categories), the inter-electrode distance (4 categories) and electrode size (3 categories)

## Results

### The effect of grid configuration on hand gesture classification accuracy

A total of 84 grid configurations were simulated with variations in the number of electrodes (7 variations: 2×2, 3×3, 4×4, 5×5, 6×6, 7×7, 8×8), distance between electrodes (4 variations: 4mm, 6mm, 8mm, 10mm) and electrode size (3 variations in radius: 1mm, 2mm, 3mm). Different configurations produced different classification accuracies, shown for a typical subject (subject 7) in Figure 3 and for all subjects in Supplementary Figure 2.

**Figure 3.**
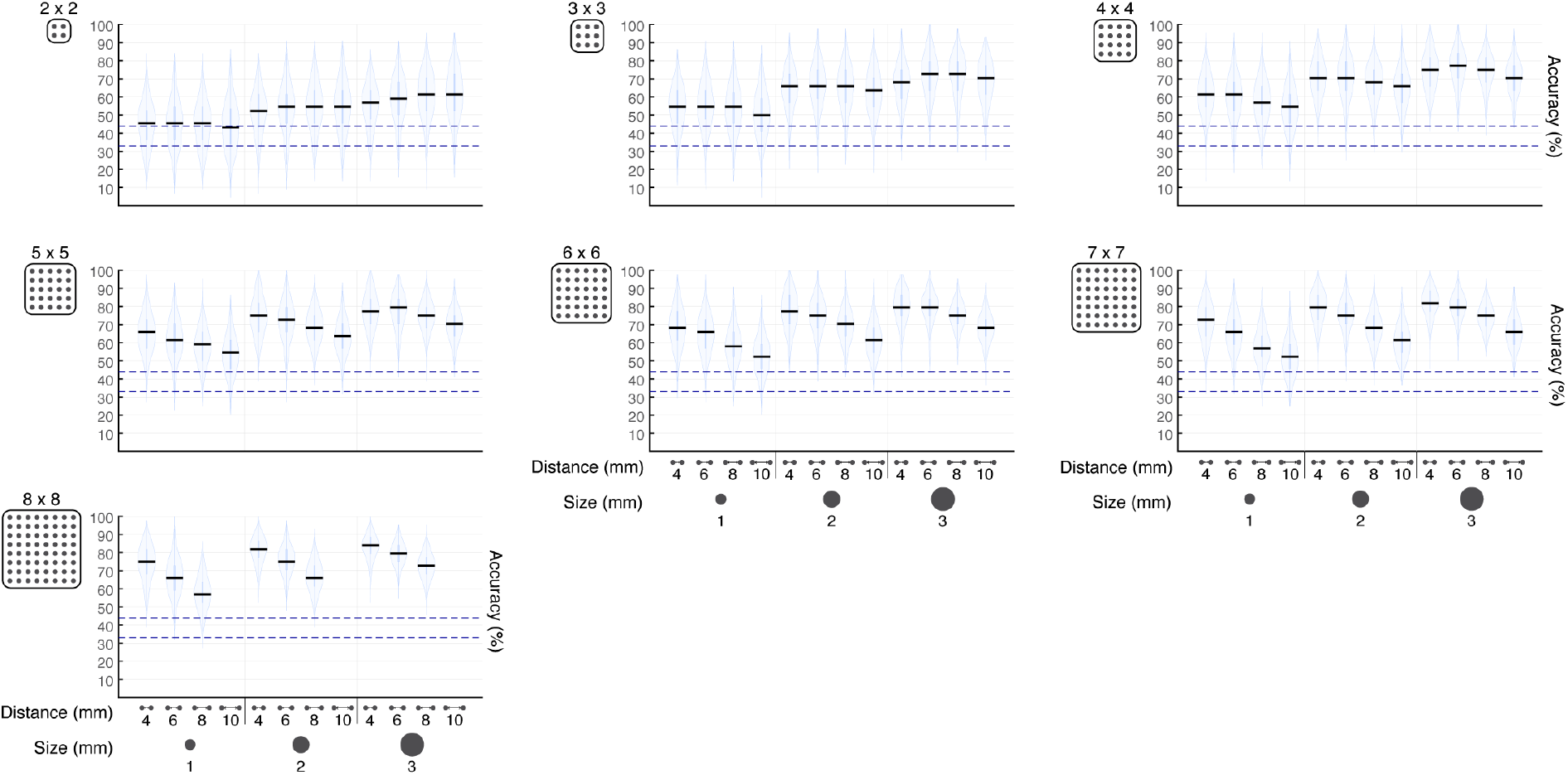
Classification accuracy as a function of grid configuration in a typical subject (subject 7). Each graph represents one variation in the number of electrodes. Each column in each graph represents a different inter-electrode distance (center-to-center) and electrode-size (radius) combination. Every violin reflects the distribution across the 2,000 scores, with a horizontal black line in every column to indicate the median. The lower blue dotted line indicates the chance level at 33%, while the upper blue line indicates the threshold (at 44%) above which the accuracy was significantly above chance.

In each individual subject, the changes in number of electrodes, distance and size showed several effects: first, more electrodes resulted in a better classification accuracy. The largest increase in classification accuracy was observed when the number of electrodes increased from smaller (e.g. 2×2) to medium sized grids (e.g. 5×5), beyond which accuracy barely improved. Second, an increase of distance between the electrodes resulted in a decrease in classification accuracy in grids with a larger number of electrodes (more than ^~^5×5). Third, increasing the electrode size resulted in a better classification accuracy in all situations. These effects were highly robust across individual subjects (Supplementary Figure 2).

The effects seen in the individual subject can also be observed in the average group classification accuracies for each of the 84 simulations (Figure 4) and the averages of the classification accuracies per significant factor (number, space, size; Figure 5). Firstly, more electrodes result in a significantly higher classification accuracy (Figure 5A, three-way ANOVA across all 10 participants: F(6, 711) = 47.42, p < .01). Secondly, an increase in distance between the electrodes significantly lowers the classification accuracy (Figure 5B, F(3, 711) = 23.81, p < .01). Thirdly, a larger electrode radius causes a significant increase in classification accuracy (Figure 5C, F(2, 711) = 162.02, p < .01). The second effect, where an increase in inter-electrode distance caused a decrease in classification accuracy, was stronger for larger numbers of electrodes, shown in a significant interaction between the number of electrodes and electrode distance (Figure 5D, F(17, 711) = 2.52, p < .01). The model accounted for a total of 47% of the variance in classification accuracy.

**Figure 4.**
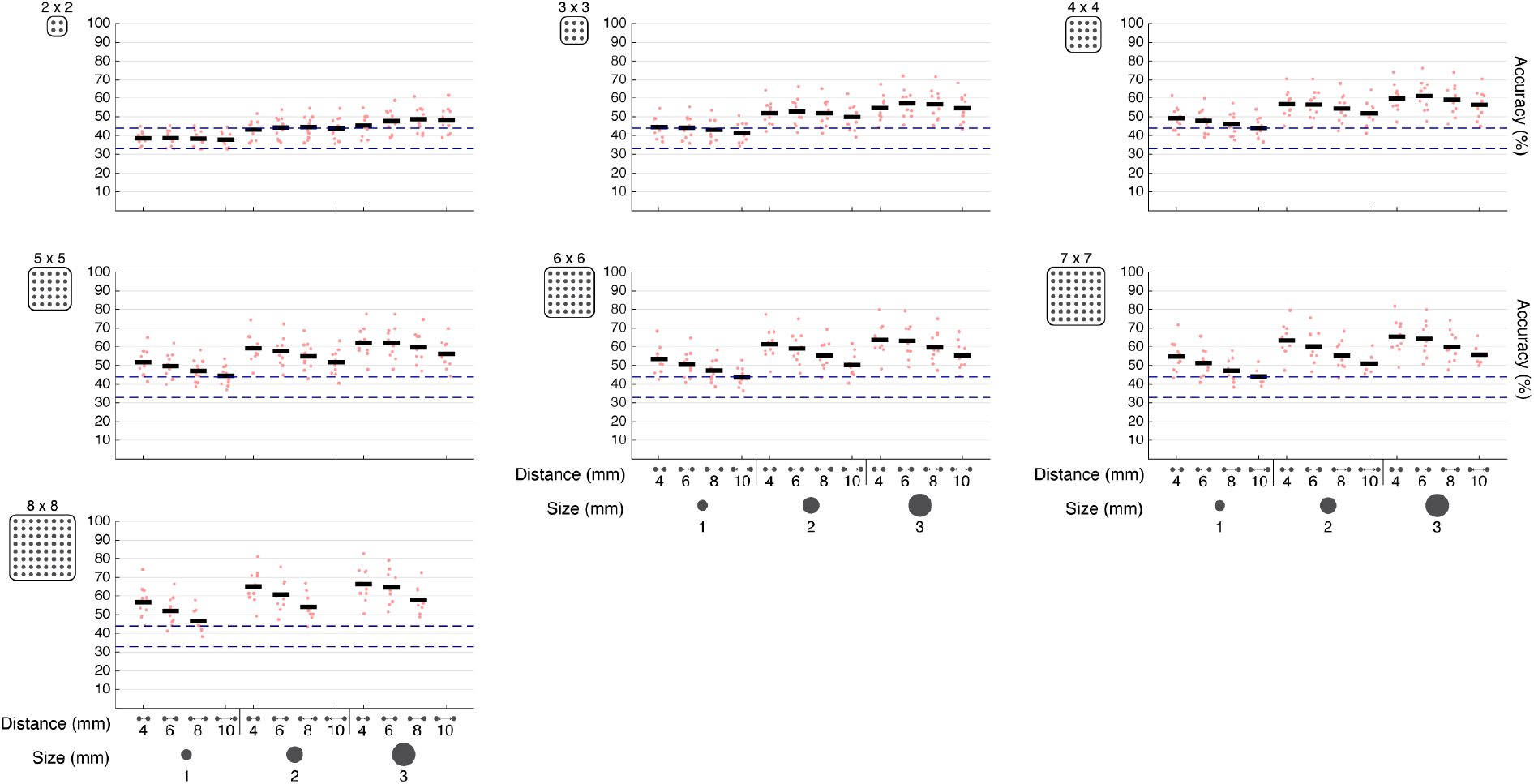
Classification accuracies for each grid configuration across the group. Each graph represents one variation in the number of electrodes. Each column in each graph represents a different inter-electrode distance (center-to-center) and electrode-size (radius) combination. The red dots indicate the average classification accuracy for every participant, with a horizontal black line in every column to indicate the average over all participants. The lower blue dotted line indicates the chance level at 33%, while the upper blue line indicates the threshold (at 44%) above which the accuracy was significantly above chance.

**Figure 5.**
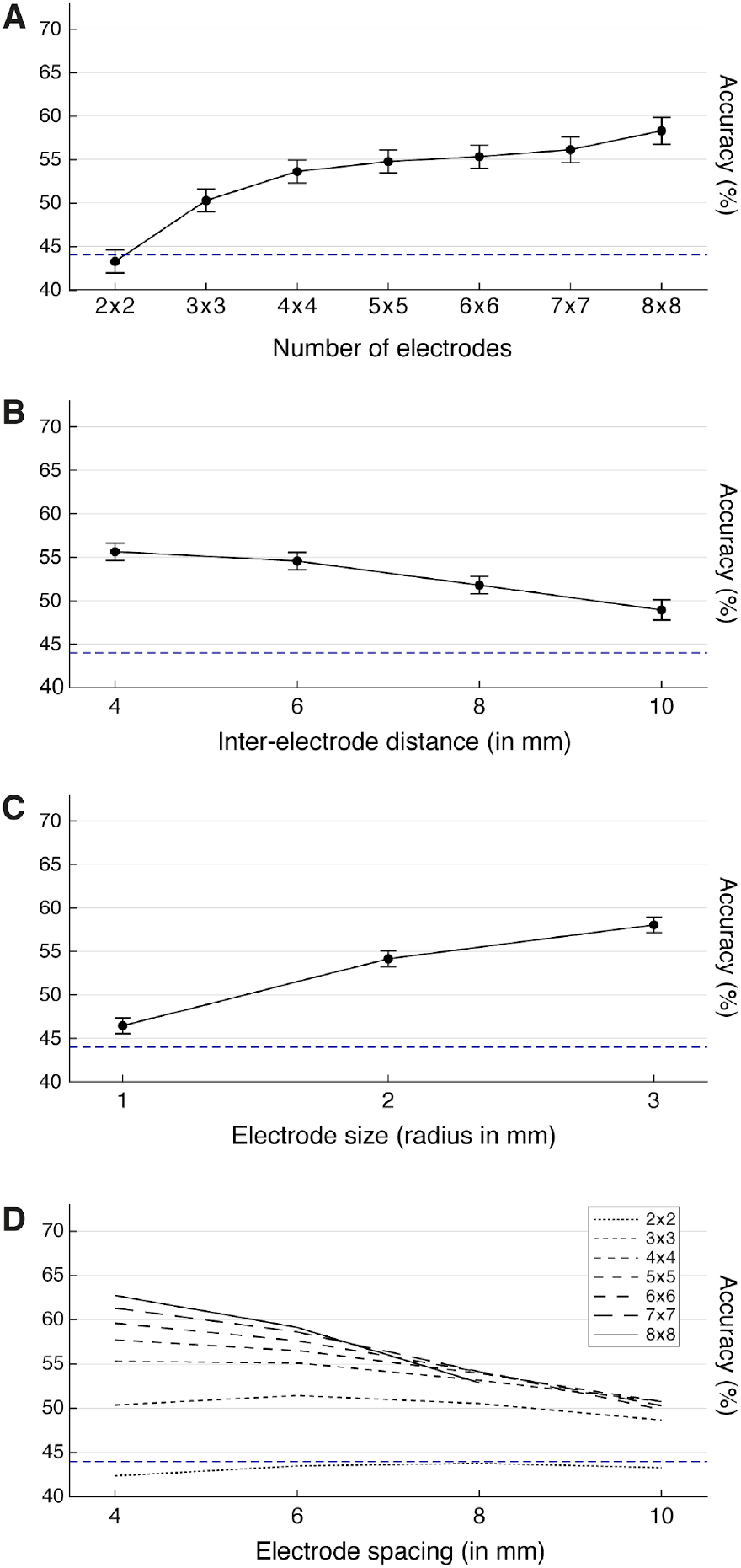
The average classification accuracy per grid configuration factor across the group. Group results showing the means for the three grid configuration parameters and the significant interaction effect. The error bars in the top three plots represent a 95% confidence interval. The blue dotted line at 44% indicates the threshold above which the accuracy was significantly above 33% chance level.

### The effect of grid configuration on hand gesture classification accuracy with placement of the entire grid within the most informative area

To investigate whether classification could be optimized by restricting all electrodes in the stimulated grids to the informative region, we ran the simulations while limiting the grids to fall only within the region of interest indicated by the searchlight (Figure 6). Some of the larger grids, with more electrodes and/or increased distances between electrodes, could not be simulated as they exceeded the region’s boundaries (Supplement Figure 1). Increasing the number of electrodes up to 5×5 electrodes on the informative region significantly increased the classification accuracy (Figure 6B, F(3, 222) = 67.43, p < .01). The effect of inter-electrode distance no longer reached significance, but became a trend (Figure 6C, F(3, 222) = 2.50, *p*=0.06) where more distance increased the classification accuracy (Figure 6). Interestingly, within this area, a larger electrode radius still significantly increased the classification accuracy (Figure 6D, F(2, 222) = 39.39, p < .01).

**Figure 6.**
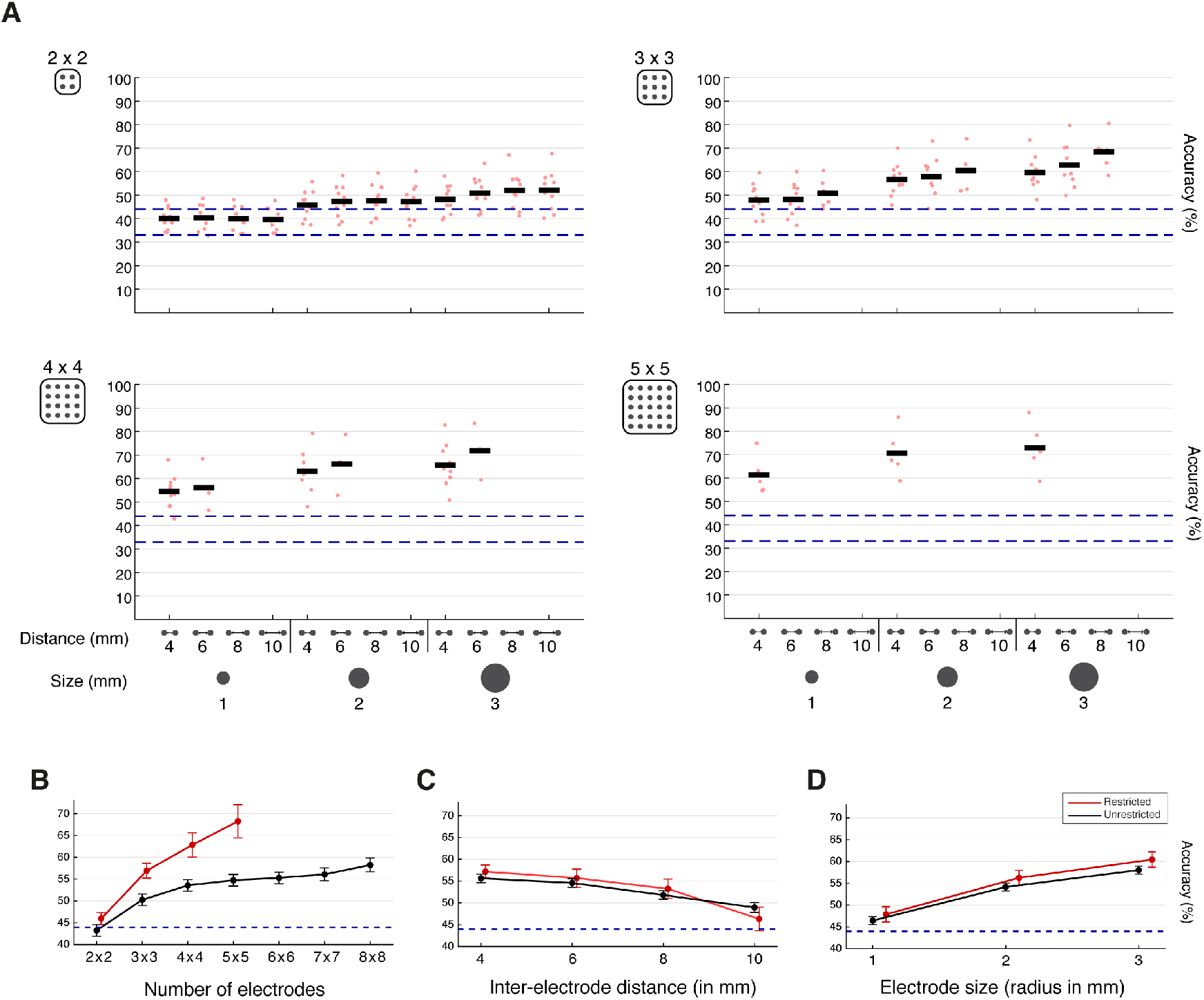
Classification accuracy as a function of grid configuration when limiting the grids to an informative region. (A) The classification accuracies on all grid configuration parameters. Each graph represents one variation in the number of electrodes, the columns represent variations in inter-electrode distance and electrode-size. Each red dot indicates a single participant, the horizontal black line indicates the average over participants. The two dotted lines indicate 33% chance level and the threshold above which the accuracy was significantly above chance at 44%. (B, C and D) The group means for the variation in number of electrodes (B), inter-electrode distance (C) and electrode size (D) are shown when the grid is placed in the informative area (red lines). For comparison, the data from Figure 5 are also displayed in black. Note that the direction of the trend in electrode space (C) might be misleading, the higher inter-electrode distances are biased to grids with a low number of electrodes, because only these grids would fit in the informative region, and therefore might result in a lower average score. The error bars represent a 95% confidence interval. The blue dotted line indicates the threshold (at 44%) above which the accuracy was significantly above 33% chance level.

Note that while the number of electrodes improved classification accuracy (Figure 6B), grids with less electrodes could also reach classification performance within the same range. Electrode grids of 5×5 electrodes, 4mm inter-distance and 3mm radius provided a classification accuracy of 73% (range 59-88) (Figure 6A bottom right); grids of 3×3 electrodes, 8mm inter-distance, 3mm radius provided a classification accuracy of 68% (range 58-80) (Figure 6A top right). Much smaller grids of 2×2 electrodes, however, showed a much lower range of classification accuracies and reached a mean accuracy of 52% (range 40-68) at 10mm inter-distance, 3mm radius.

### Optimal grid placements can be identified to improve hand gesture classification performance

The simulation results may be used to predict grid configurations with high classification accuracy in individual subjects for BCI purposes. In addition, some grids can exceed the average classification accuracy of that configuration when positioned in the correct way. As a proof of concept, we performed an agglomerative cluster analysis on the 2,000 grid placements within three specific grid configurations (3×3, 8mm inter-distance, 3mm radius; 4×4, 8mm inter-distance, 2mm radius; 5×5, 4mm inter-distance, 1mm radius). Within each configuration, the 2,000 grids were spatially clustered based on the 3D positions of two corner electrodes, while limiting the euclidean clustering distance to 2mm. This results in clusters of grids that have similar position and angle. Figure 7 shows, for three of the grid configurations, the cluster that provided the highest average classification accuracy (over the grids in that cluster) for a single subject (subject 7). Grid placement suggestions for all participants are shown in Supplementary figure 3. We required clusters to have at least 3 grids. As demonstrated in Supplementary figure 4, different euclidean clustering distances will provide different clusters, which consist of different numbers of grids. The clustering method and parameters we have used here for this proof of concept is just one of the ways to find the optimal position based on the simulated data. In terms of grid placement, the simulated grids and clustering can be used to find a balance between the spatial margin of error in placement (i.e. the number of grids in the simulation and their clustering distance) and the expected classification performance.

**Figure 7.**
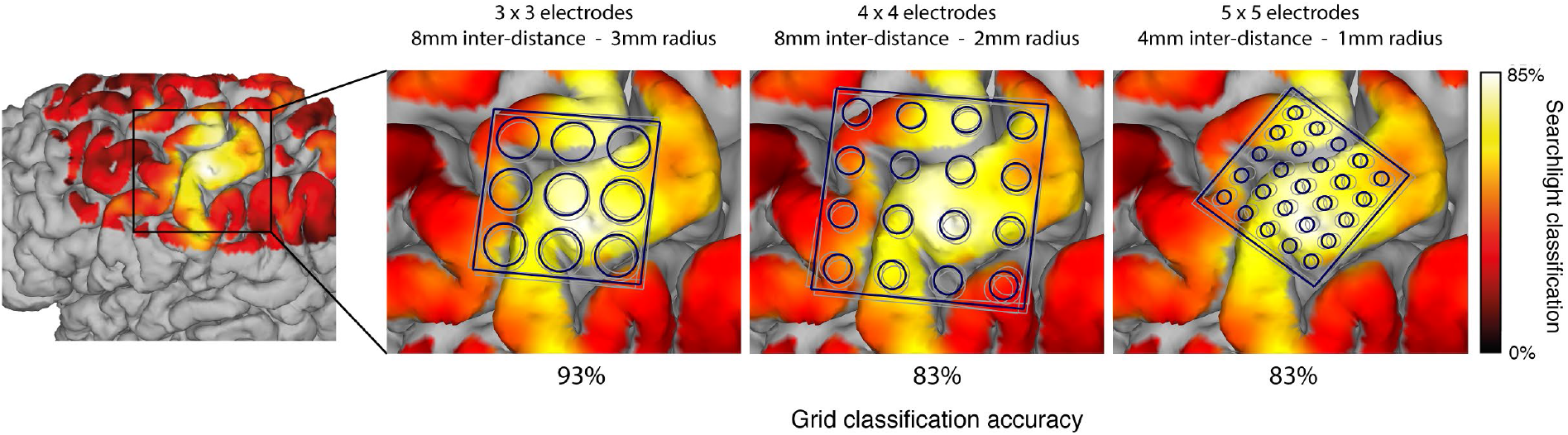
Proof of concept for the optimal placement of three different grid configurations in subject 7. The leftmost image shows a heat-map of the brain that indicates the presence of hand-gesture information based on searchlight classification. The three images on the right show an optimal placement within three different grid configurations. Each of the suggestions is based on an agglomerative cluster analysis on the 3D positions of two corner electrodes of the 2,000 grid placements within that grid configuration. Grids, limited to an euclidean distance of 2mm, were clustered together. The grid-cluster with the highest average classification accuracy (over all grids) is shown for each of the three grid configurations, the individual grids in that cluster are shown in grey and their average position in blue. The average classification accuracies over the grids in the cluster is shown underneath the image.

## Discussion

This study investigated how different simulated ECoG grid configurations capture information on executed hand-gestures from the brain surface. To this end, 7T fMRI data were used as a proxy for neural activity, allowing for virtual placement of a range of grid configurations and parameterized assessment of electrode number, size and distance. Our simulations show that increasing the number of electrodes improves hand-gesture decoding accuracy, with the largest improvement seen up to 5×5 electrodes. Reducing inter-electrode distance and increasing electrode size (to 3mm in radius) improved hand gesture decoding accuracy.

Decoding accuracy was further optimized when grids were restricted to an informative region for hand-gestures as identified in individual subjects by searchlight classification. We found good classification scores (68%, with chance at 33%) for grids with 3×3 electrodes, 8mm inter-electrode distance and a 3mm electrode radius. Adding electrodes up to 5×5 at 4mm inter-distance and 3mm size provided small increases in classification accuracy (73%). These grid resolutions are commensurate with the reported resolution of finger representations in human sensorimotor areas. Within the sensorimotor hand area, detailed finger representations have been demonstrated with electrical stimulation (Penfield and Boldrey 1937), fMRI (Beisteiner et al. 2001; Hlustik et al. 2001; Dechent and Frahm 2003; Olman et al. 2012; Schellekens et al. 2018) and ECoG (Miller et al. 2009; Siero et al. 2014). Within these sensorimotor hand areas, clinical ECoG grids with a standard resolution (mostly 4×8 to 8×8 electrodes, spaced 10mm center-to-center with ^~^2.3mm exposed surface diameter) have been used to differentiate between finger representations (Miller et al. 2009), decode separate finger movements (Shenoy et al. 2007; Kubánek et al. 2009), decode five distinct hand postures (Chestek et al. 2013), two different grasp types (Pistohl et al. 2012), decode reaching movements (Bundy et al. 2016) and simple hand/elbow movements (Yanagisawa et al. 2012) using the changes in the high frequency band. Siero et al. (Siero et al. 2014) found that the activation foci of the different fingers (thumb to little finger) span ^~^10mm in both fMRI and ECoG, with an estimated distance of 3-4 mm between thumb, index and little finger. Dechent (Dechent and Frahm 2003) found a similar span in the center-of-mass distance estimates in fMRI, ranging from 6.6mm to 16.8mm. With regard to the distance between finger representations, the distance between centers of mass does not necessarily mean those yield optimal decoding performance, since overlapping finger representations between centers (Schellekens et al. 2018) may not affect center of mass location but it does affect discriminability of separate finger movements. As a result, one may expect optimal distance to be larger for decoding than for mapping. A study that decoded facial expressions from the sensorimotor cortex showed that only part of a high-density grid surface (15×15mm to 20×20mm based on 6×6 electrodes having 3mm or 4mm inter-electrode distance) was enough to provide good decoding accuracy (Salari et al. 2020). This supports the idea that a limited number of electrodes with optimal inter-electrode distance on the most essential area is enough to provide high decoding performance.

Grids with more electrodes at denser inter-electrode distances (e.g. 5×5 electrodes, 4mm distance, 3mm size) provided only small increases in classification accuracies in comparison to smaller grids with a higher inter-electrode distance (e.g. 3×3 electrodes with 8mm distance, 3mm size). Similar to ECoG studies with standard grids, studies using high-density grids on sensorimotor cortex (4×8 and 8×8 electrodes, distanced 3mm apart with 1.3mm exposed surface diameter) were able to distinguish finger representations (Siero et al. 2014) and decode four hand-gestures (Bleichner et al. 2016; Branco et al. 2017) using HFB changes. However, it remains difficult to directly compare the performance between low and high-density grids in individual subjects. We can only speculate on how an even denser grid would perform. A spatio-spectral analysis using 64 electrode-strips with an inter-electrode distance of 0.5mm indicated an optimal distance of 1.25mm (Freeman et al. 2000). A study combining spatio-spectral analyses from rat recordings and a dipole model suggested a minimal spacing of 1.7–1.8 mm for subdural and 9–13 mm for epidural grids to minimize signal overlap between electrodes (Slutzky et al. 2010). However, current levels of fMRI voxel resolution are not likely to be accurate enough to simulate such electrode distances, given a certain point-spread due to non-discrete anatomical properties of the microvasculature that gives rise to the BOLD effect. Still, the decoding of more gestures, movements, or more complex behaviors (Chang et al. 2010; Kellis et al. 2010; Anumanchipalli et al. 2019; Salari et al. 2019) may benefit from denser grids. The methods we share here can be used by others to evaluate the decoding of more complex behaviors for BCI purposes.

Larger electrodes (up to 3mm in radius) decoded better. There are two aspects that may play a role in this optimal simulated size: the averaging of fMRI signals across voxels, and the averaging across neuronal populations. First, fMRI BOLD signals contain noise that is reduced by averaging across voxels (Edelstein et al. 1986; Parrish et al. 2000). In our simulation, the electrodes with a larger radius included more underlying voxels. For classification, each virtual electrode served as an input feature to the classifier based on the average signal of all the voxels underneath the electrode. Therefore, larger electrodes average over a higher number of voxels, increasing the signal-to-noise ratio of every feature in the classifier. The increase in signal-to-noise ratio has likely contributed to classification accuracy. Second, field potential studies have shown that sampling neurophysiological signals from a larger patch of cortex can be highly informative (Toda et al. 2011; Ibayashi et al. 2018; Kanth and Ray 2020). A recent study (Kanth and Ray 2020) showed that ECOG recordings can provide more relevant information and higher decodability than microelectrode recordings. ECoG electrodes could effectively average over a neural population, negating the noise in the signal while preserving the common relevant information. Since the BOLD signal in a voxel is a representation of a neural population, the increase in classification accuracy that comes with larger electrode size (i.e. more voxels) might not be solely due to fMRI specific vascular noise, but could also be attributed to a reduction of noise by sampling from a larger neuronal population.

Although the fMRI BOLD signal is a more indirect measure of neuronal activity, it may serve as a good approximation of ECoG measurements. Several studies have shown that electrophysiology and BOLD signal changes correlate across time (Logothetis et al. 2001; Mukamel et al. 2005), that HFB power and BOLD correlate spatially with matching peak activity (Conner et al. 2011; Hermes et al. 2012, 2014), that the non-linearities observed in fMRI can be predicted by ECoG HFB signal (Siero et al. 2013) and that the level of BOLD increase matches the level of ECOG HFB power increase across conditions (Winawer et al. 2013; Jacques et al. 2016; Hermes et al. 2017). One study (Siero et al. 2014) specifically combined fMRI measurements (voxel size of ^~^1.5mm) and high-density ECoG measurements (8×4 electrodes, spaced 3mm with a diameter of 1mm) and found distinguishable finger representations that matched between the two techniques. The simulations based on the BOLD responses are likely to be indicative for what would be found with ECOG measurements.

The optimal grid resolution and placement is important for intracranial BCI applications that target patient populations. While our study included healthy participants, previous research suggests that hand representations are maintained in patients with an amputated arm (Bruurmijn et al. 2017). Moreover, ECoG based BCI applications can benefit from fMRI localizer tasks to provide non-invasive identification of the informative region prior to surgical implantation (Vansteensel et al. 2016). Several BCI systems in patients with ALS or paraplegia have used fMRI to place grids on sensorimotor cortex (Collinger et al. 2013; Bouton et al. 2016; Vansteensel et al. 2016; Benabid et al. 2019) and visual cortex (Beauchamp et al. 2020). While fMRI is normally used only to identify the relevant region, the method described in this article can also advise on the optimal placement and configuration of a grid.

When an ECoG grid is implanted for BCI purposes, it is important to consider a reduced number of electrodes, given practical considerations such as the cost and complexity of high channel-count implantable amplifiers and real time signal processing (Leuthardt et al. 2009; Reisch et al. 2013). In addition, the surgical procedure and implant could potentially carry a risk to the patient. Implanting a larger grid requires a larger surgical exposure, which may impose a slight risk for complication (Wong et al. 2009). As our results suggest, the grid should be focused on the region of the brain that contains the most hand-gesture information. Our results provide a principled way of computing the tradeoff between grid size, density and decoding performance. The pipeline that we used for simulation could be used pre-surgically to locate the most informative area, find the grid configuration that would lead to the best performance and suggest one or more optimal positions on top of the most informative area for individual patients.

## Conclusion

Most of the information encoding hand gestures is densely packed within a small area of the sensorimotor cortex. Our results show that optimal decoding of hand-gestures is achieved by placing a grid within this informative region. Grids of 3×3 electrodes with an inter-electrode distance of 8mm and electrode size radius of 3mm provided good classification accuracy. It has been assumed that densely-packed, small diameter, electrodes will provide the best resolution of functional representation. Based upon this fMRI-modeling approach, we might reassess this assumption. These densely spaced configurations might provide only a marginal benefit while being less practical in terms of complexity and cost of high channel-count implantable amplifiers, real-time processing and clinical invasiveness. As one might expect, positioning of the electrodes on the most informative area has a strong influence on the classification accuracy. The simulation techniques outlined in this article can be used more generally in clinical and research settings to identify optimal grid configurations and placements for neural prosthetics.

## Acknowledgements

We are grateful for the participation of the volunteers in the study at UMC Utrecht. We would like to thank Melissa Koppeschaar for her assistance in the data collection, Philippe Cornelisse for operating the MRI scanner, Matthijs Raemaekers for his help in correcting the geometric distortions in the functional data, Mariana Branco for putting us on track of the NeuralAct (cortex hull generation) routines and John Burkardt’s (3D intersection) routines, and Mark Bruurmijn for the development of the task.

This work was supported by the NIH-NIMH CRCNS R01MH122258-01 (DH, KJM, MvdB), the NIH-NCATS CTSA KL2TR002379 (KJM) and ERC-Adv 320708 (NR, MvdB). The funding source did not have any role in study design; in the collection, analysis and interpretation of data; in the writing of the report; and in the decision to submit the paper for publication. Manuscript contents are solely the responsibility of the authors and do not necessarily represent the official views of the NIH. KJM is also supported by the Van Wagenen Foundation and by the Brain Research Foundation, with a Fay/Frank Seed Grant.

## Supplementary material

**Supplementary Figure 1.**
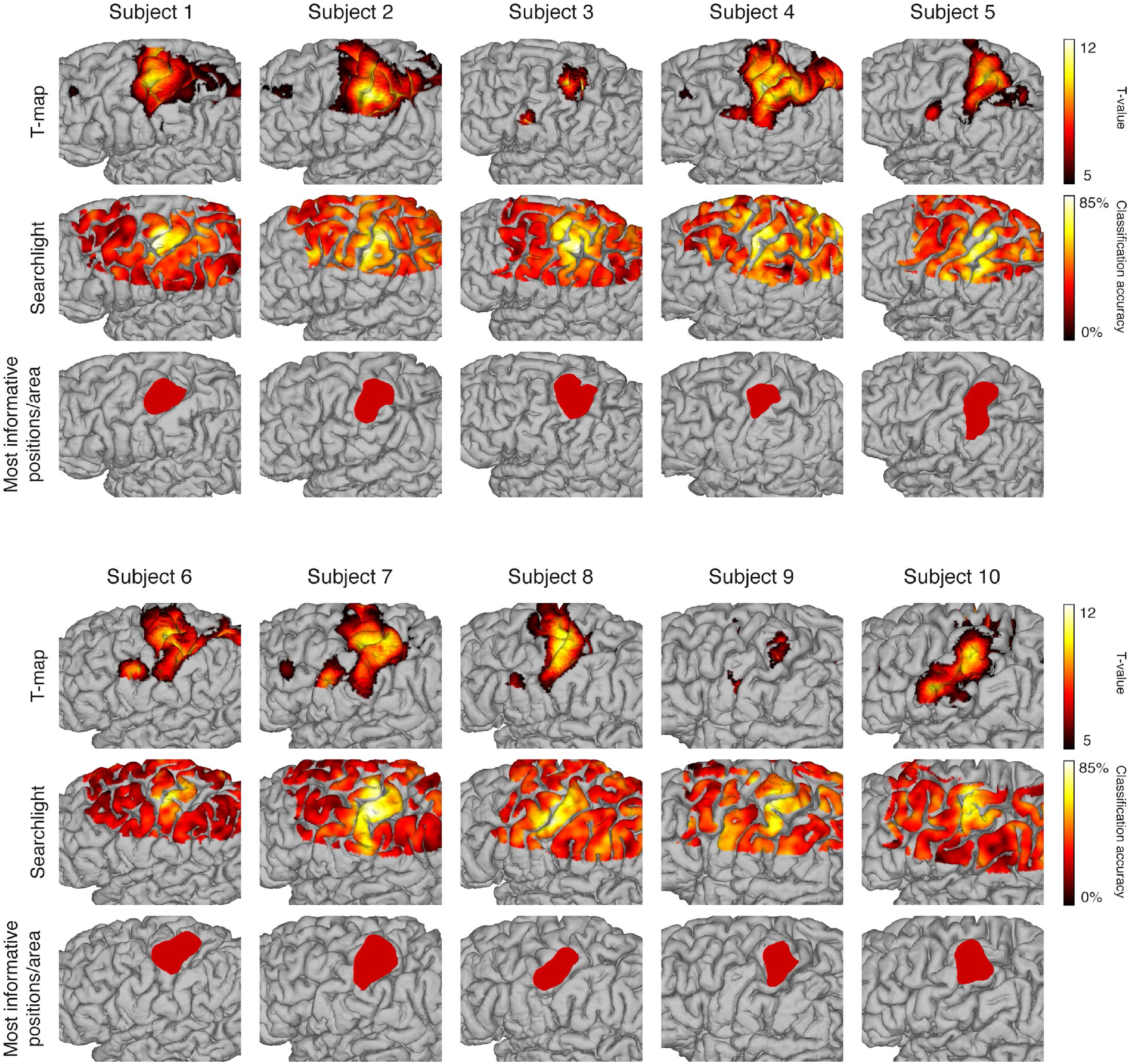
fMRI BOLD activity, information-mapping and most informative region during performance of hand-gestures per participant. The top row shows significant activation (based on a GLM with a single regressor, gesture vs rest, Bonferroni corrected). The middle row shows the averaged classification result of 200,000 searchlights that were placed at random dorsolaterally over the frontal and parietal lobes. The bottom row shows only the positions of the searchlights with a classification accuracy above the 95th percentile, limited to the largest cluster. Together, these positions empirically identify the cortical hand-gesture area and were used as the centers for the placement of the grids that were simulated.

**Supplementary Figure 2.**
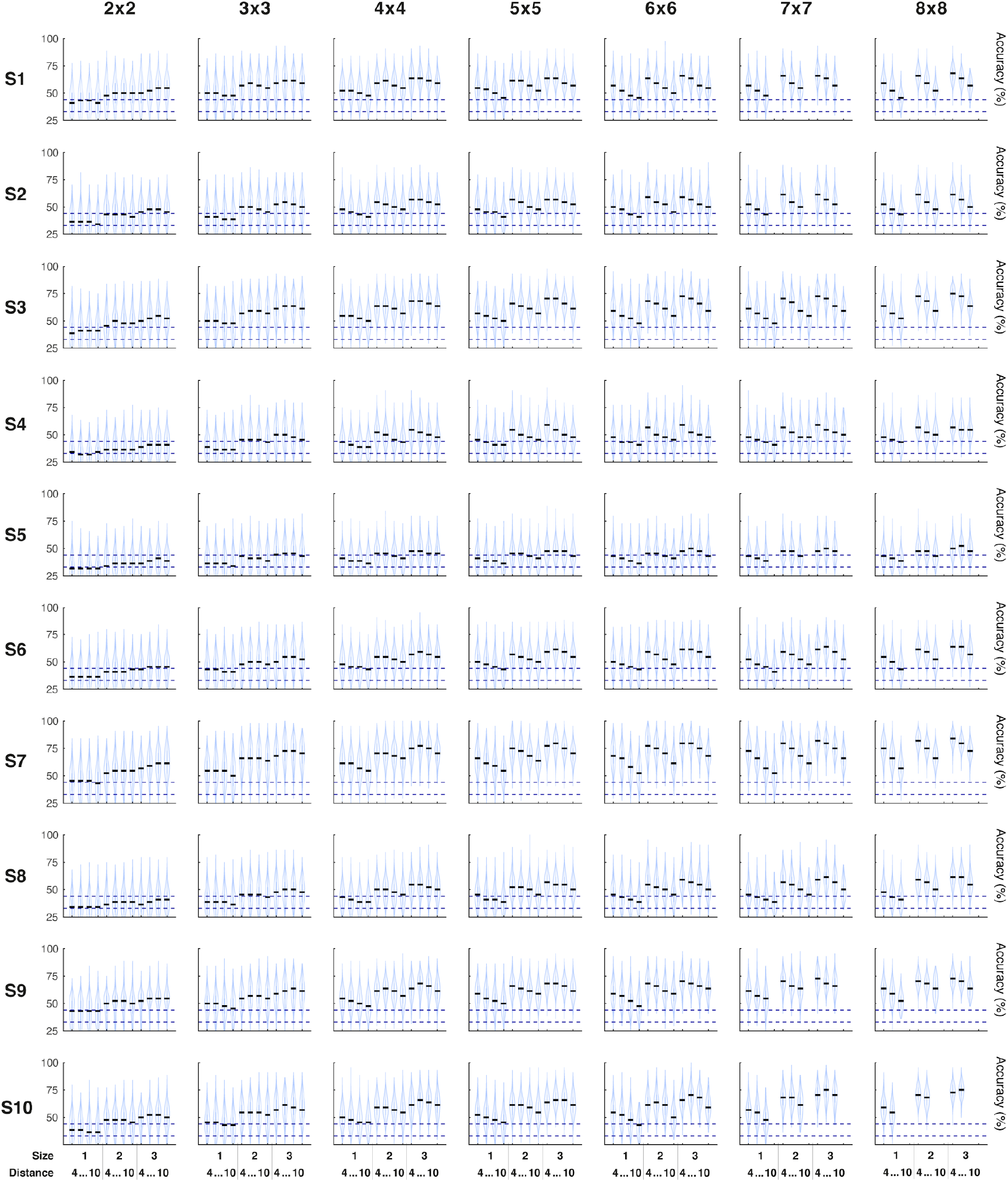
Violin plots showing the classification accuracy scores for all grid configurations in all subjects. Placement was not limited to the most informative region. Rows represent the 10 subjects, columns the number of electrodes. Each column in each graph represents a different inter-electrode distance (4mm, 6mm, 8mm, 10mm) and electrode-size (1mm, 2mm, 3mm) combination. The violin reflects the distribution of scores, with a horizontal black line in every column to indicate the median. The lower blue dotted line indicates the chance level at 33%, while the upper blue line indicates the threshold (at 44%) above which an accuracy score is significantly above chance.

**Supplementary Figure 3.**
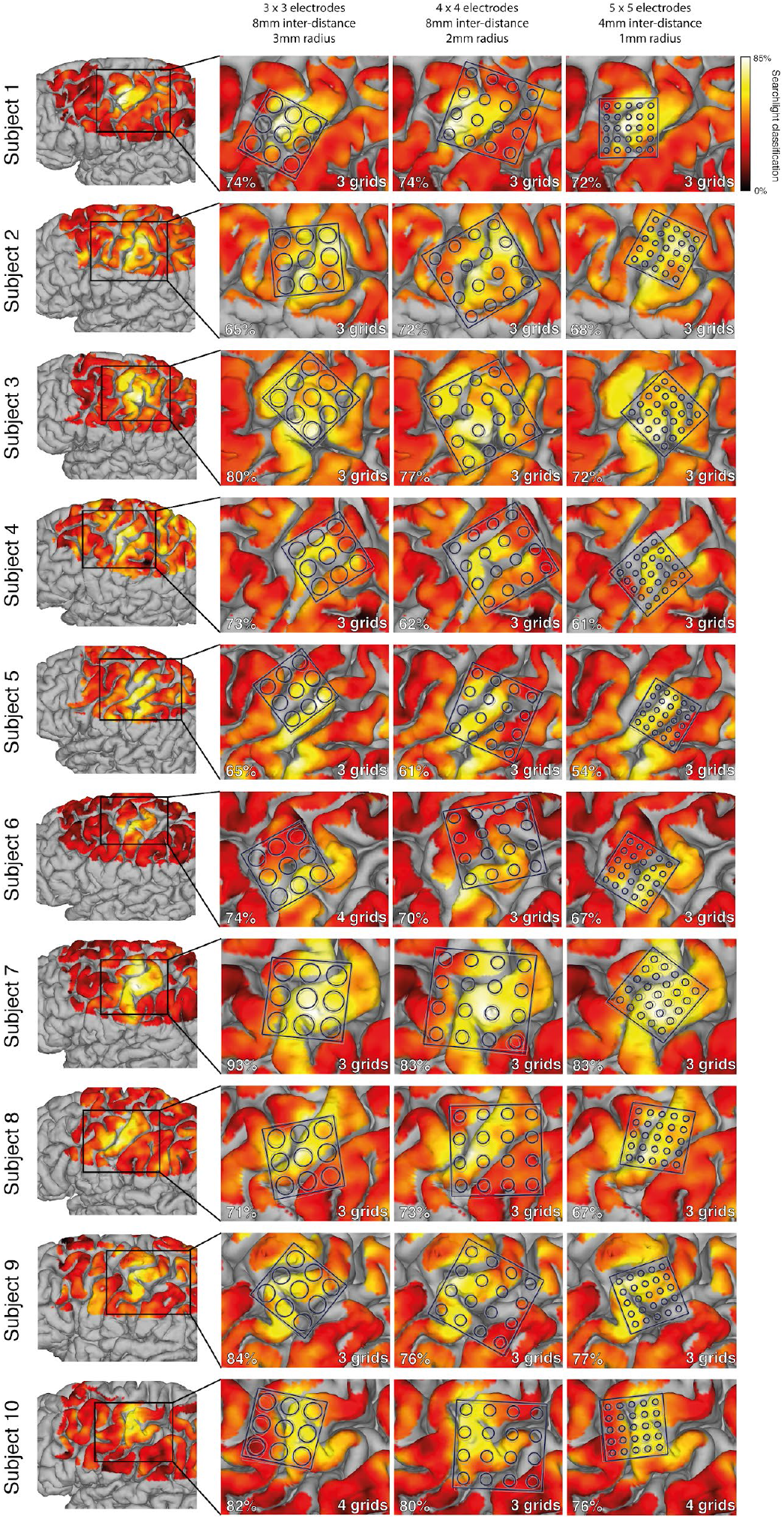
Suggestions for the placement of three different grid configurations in all subjects. Each row represents one subject. The first column shows the presence of hand-gesture information over the brain based on searchlight classification. The other three columns suggest an optimal placement of three different grid configurations. Each of the suggestions is based on an agglomerative cluster analysis on the 3D positions of two corner electrodes of the 2,000 grid placements within a grid configuration. Grids were clustered together by euclidean distance with a cutoff distance of 2mm. Each image shows the grid-cluster that - within one grid configuration for a specific subject - had the highest average classification accuracy. The individual grids in that cluster are shown in grey and their average position in blue. The average classification accuracies over the grids in the cluster are shown in the bottom-left of each image, the number of grids that make up a cluster is noted on the bottom-right.

**Supplementary Figure 4.**
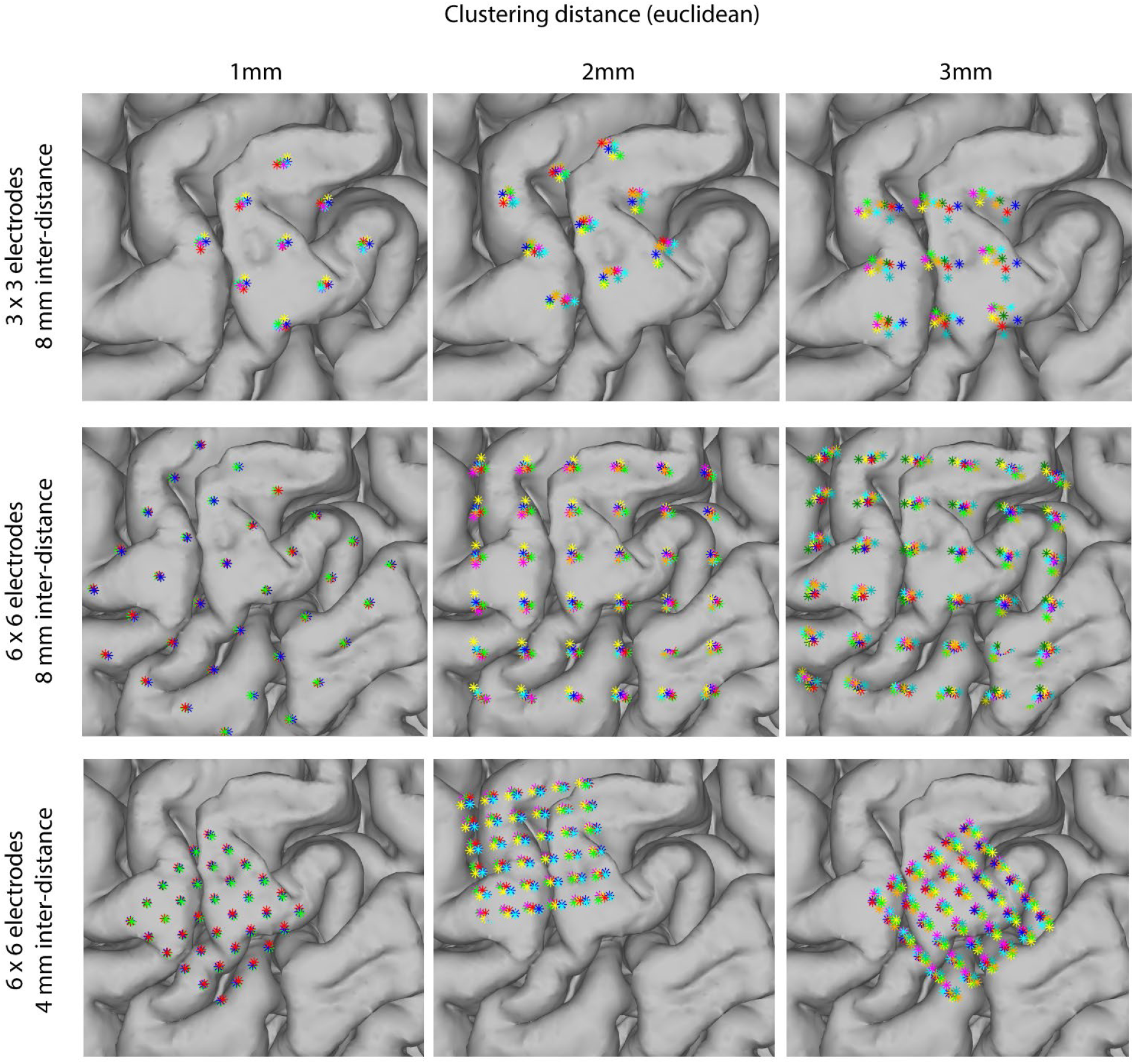
Different euclidean clustering distances, number of electrodes and inter-electrode distances result in different clusters. The rows represent three different grid configurations that vary in number of electrodes and inter-electrode distance, for subject 10. The columns represent the three euclidean distance clustering parameters (1mm, 2mm and 3mm). For each image, the cluster with the highest number of grids was selected; each color represents a single grid in that cluster. The images illustrate how the resulting cluster will differ depending on the clustering method and parameters in combination with the grid configuration. Choices should be made, taking into account the margin of error in placement and the predicted classification performance.

